# Exploring Motor Network Connectivity in State-Dependent Transcranial Magnetic Stimulation: A Proof-of-Concept Study

**DOI:** 10.1101/2023.06.29.547027

**Authors:** Laura Marzetti, Alessio Basti, Roberto Guidotti, Antonello Baldassarre, Johanna Metsomaa, Christoph Zrenner, Antea D’Andrea, Saeed Makkinayeri, Giulia Pieramico, Risto J. Ilmoniemi, Ulf Ziemann, Gian Luca Romani, Vittorio Pizzella

## Abstract

State-dependent non-invasive brain stimulation (NIBS) informed by electroencephalography (EEG) has contributed to the understanding of NIBS inter-subject and inter-session variability. While these approaches focused on local EEG characteristics, it is acknowledged that the brain exhibits an intrinsic long-range dynamic organization in networks.

This proof-of-concept study explores whether EEG connectivity of the primary motor cortex (M1) in the pre-stimulation period aligns with the motor network (MN) and how MN state affects responses to transcranial magnetic stimulation (TMS) of M1. One thousand suprathreshold TMS pulses were delivered to left M1 in 8 subjects at rest, with simultaneous EEG. Motor evoked potentials (MEPs) were measured from the right hand. Source-space functional connectivity of left M1 to the whole-brain was assessed using the imaginary part of the Phase Locking Value at the frequency of the sensorimotor µ-rhythm in a 1-second window before the pulse. Group-level connectivity revealed functional links between left M1, left supplementary motor area, and right M1. Also, pulses delivered at high MN connectivity states result in a greater MEP amplitude compared to low connectivity states. At single-subject level, this relation is more expressed in subjects that feature an overall high cortico-spinal excitability. In conclusion, this study paves the way for MN connectivity based NIBS.

**Highlights:**

- EEG pre-stimulus connectivity of left M1 largely corresponds to the motor network
- Stronger motor network (MN) connectivity corresponds to greater MEP amplitudes
- Linear regression models based on MN connectivity predicts MEP amplitude

## 1. Introduction

For over three decades, non-invasive brain stimulation (NIBS) has been used to modulate brain activity in healthy subjects and patients (Hallett, 2007, 2000; Rossini et al.; 2015), for scientific, diagnostic and therapeutic purposes. However, effects are highly variable, thus limiting its clinical use (Ziemann and Siebner, 2015). Recently, it has become clear that, to reduce the variability of the stimulation effects not only between subjects but also between sessions, the internal state of the brain before NIBS must be taken into account (Bergmann, 2018; Silvanto et al.; 2008). For this purpose, the integration of NIBS with techniques able to non-invasively measure neuronal activity, such as electroencephalography (EEG; (Bergmann et al.; 2016; Ilmoniemi and Kičić, 2010)), has offered a window into the state of the brain before the stimulation (Bai et al.; 2022; Zrenner et al.; 2018). This brain state has largely been assessed by looking at the spectral characteristics of the EEG signal at the channels nearby the stimulation site (Mäki and Ilmoniemi, 2010). In particular, the phase of the sensorimotor 9-13 Hz µ-rhythm has been considered as an indicator of cortical excitability that determines the response to Transcranial Magnetic Stimulation (TMS; Desideri et al.; 2019; Schaworonkow et al.; 2019; Zrenner et al.; 2018), although the observed phase effects vary with varying stimulation and analysis parameters (Karabanov et al.; 2021; Madsen et al.; 2019).

In parallel with advances in EEG-informed NIBS, neuroscience has seen a paradigm shift from a modular view, in which different functional units act as independent processors, to a large-scale network view, in which dynamic interactions between brain areas are crucial for cognition and behavior (Bressler and Menon, 2010). While functional magnetic resonance studies have been seminal in this regard (Fox and Raichle, 2007), non-invasive electrophysiology has contributed to this view by the characterization of neuronal networks in terms of their oscillatory fingerprints (Brookes et al.; 2011; de Pasquale et al.; 2010; Engel et al.; 2013; Ganzetti and Mantini, 2013; Mantini et al.; 2007; Marzetti et al.; 2019, 2013), a view largely supported by the Communication Through Coherence (CTC) hypothesis (Fries, 2015, 2005). In this framework, a brain state can be described as the evolving dynamics of one or more large-scale networks (Kringelbach and Deco, 2020), including the so-called resting-state networks (Deco and Corbetta, 2011), that constrain ongoing activity in the absence of any externally imposed task. Several studies have investigated the effects of invasive and non-invasive stimulation on resting-state networks and, more in general, on remote regions connected to the stimulation site (Beynel et al.; 2020; Boutet et al.; 2019; Pieramico et al.; 2023). However, while the techniques for connectomic neuromodulation studies appear mature (Horn and Fox, 2020; Ozdemir et al.; 2020), scarce evidence has been provided, so far, for the impact of network dynamics on stimulation effects, explicitly using functional connectivity approaches (Ferreri et al.; 2014). In addition, so far only one study (Vetter et al.; 2023) has investigated sensor level functional connectivity as a feature for brain-state dependent stimulation, although the accuracy in space and time is limited by the sensor-level analysis and by the real-time software implementation.

The aim of this work is to bridge the gap between the study of brain networks with non-invasive electrophysiology and brain state-dependent stimulation, with the long term goal of systematically using EEG-derived brain networks to drive the stimulation in space and time with millimeters and milliseconds resolution (Marzetti et al.; 2024; Sinisalo and Rissanen et al.; 2024). Here, we provide a proof-of-concept that fast-dynamic brain networks (Marzetti et al.; 2024) can be derived from combined EEG-TMS data in the pre-stimulation resting period and that the momentary connectivity state of such networks is related to the stimulation end-point. Specifically, our proof-of-concept assessed, using data from a previous study (Metsooma et al.; 2021) and the phase-locking of the oscillatory µ-rhythm, the functional connectivity between the primary motor cortex (M1) signal and the signals at all other brain locations, and its putative relation to the amplitude of the Motor Evoked Potentials (MEPs). The choice of the µ-rhythm frequency for our analysis is driven by the observation that ongoing oscillations in the Motor Network (MN) at rest are expected to synchronize at this frequency (Hari, 2006). Specifically, we hypothesized that: 1) the pattern of long-range functional connectivity of the primary motor cortex (M1) in the pre-stimulation period largely overlaps the spatial topography of the MN; 2) the connectivity state of this network impacts the effect of TMS pulses delivered at M1 on a trial-by-trial basis; 3) such impact is augmented if not only connectivity properties but also local properties are considered.

## 2. Material and methods

### 2.1 Participants and experiment

Eight right-handed adults (5 female, 3 male; mean±SD age 23.5±3.3 years) with no history of neurological and/or psychiatric pathologies were enrolled and correctly completed the study. All participants gave written informed consent before participation. The study was approved by the local ethics committee at the University of Tübingen and conducted in accordance with the Declaration of Helsinki. Data were acquired at the University of Tübingen using a concurrent EEG−TMS setup in a single session for each participant (duration about 3 hours). EEG and electromyography (EMG) were simultaneously recorded (sampling rate 5 kHz). EEG was recorded using a TMS-compatible 128-channel cap (EasyCap BC-TMS-128, EasyCap, Herrsching, Germany) positioned according to the International 10–5 system. EMG was recorded from the abductor pollicis brevis (APB) and first dorsal interosseous (FDI) muscles of the right hand in a bipolar belly-tendon montage. A TMS stimulator (PowerMAG Research 100, MAG & More, Munich, Germany) was used to deliver biphasic pulses through a figure-of-eight coil (PMD70-pCool, 70-mm winding diameter, MAG & More, Munich, Germany). The relative head and coil positions were tracked using optical neuronavigation (Localite GmbH). After preparation of EEG, EMG, neuronavigation and pinpointing the EEG electrodes, the hand representation of left M1 (lM1) was targeted orienting the coil such that the strongest field was induced in a posterior-lateral to anterior-medial direction. The motor hotspot was defined as the position and orientation of the coil requiring the smallest stimulation intensity to evoke MEPs in either of the two hand muscles. Resting Motor Threshold (rMT) was defined as the minimum stimulation intensity able to elicit MEPs with peak-to-peak amplitudes > 50 µV in 50% of test pulses (Groppa et al.; 2012). During the experiment, participants were seated comfortably and fixated a cross located approximately 1 m in front of them. One thousand single TMS biphasic pulses were then applied, in a single session with an interstimulus interval of 2±0.25 s at a stimulation intensity of 110% rMT. A different analysis using the same dataset has been previously reported (Metsomaa et al.; 2021).

### 2.2 Data processing

EEG data were downsampled to 1 kHz and split into windows ranging from –1004 to –4 ms relative to the TMS pulse. A Laplacian-based trend detection was applied to remove slow trends in the data, and noisy or bad channels/trials were identified and removed. A subject-level independent component analysis (ICA) was then performed using the FastICA technique (Hyvarinen, 1999) in the subspace generated by the 35 largest principal vectors. Further details are given in (Metsomaa et al.; 2021) and in the preprocessing source code at www.github.com/bnplab/causaldecoding.

For each independent component (IC), the channel-level topographies and power spectra were calculated and visually inspected by two experienced researchers; ICs with a clear artifactual hallmark were discarded. The remaining ICs were projected to the source-space using the eLORETA spatial filter (Pascual-Marqui et al.; 2011) to identify the corresponding neural generators. ICs whose source-space topography showed maxima over the motor cortices were considered for the identification of the individual µ-rhythm frequency, defined as the frequency in the 9–13-Hz range at which a maximum in IC power spectrum occurs. Among all ICs, only those with a clear peak at the µ frequency and a motor signature in source space were used to reconstruct channel-level cleaned signals for further analysis. This step allowed us to disentangle the contribution of µ-rhythm activity from the original signal.

EMG data were divided into stimulus-locked trials ranging from –500s to 500 ms relative to the TMS pulse. First, slow drifts were removed by trendline fitting; then, 50-Hz noise was removed (Metsomaa et al.; 2021). Visual inspection of EMG data allowed to remove bad trials, such as those presenting clear EMG activity before the pulse. TMS-related artifacts were removed by relying on an exponential fitting. For each trial and channel, the peak-to-peak MEP amplitude was estimated as the EMG signal range of the manually defined window of data that contains the MEP (Metsomaa et al.; 2021). Finally, an across-trials principal component analysis was applied to the log-transformed MEP amplitudes estimated from the FDI and APB muscles; the first principal component was used in the subsequent analyses as it explained 99.3±0.5% of the variance (mean and standard deviation across subjects).

### 2.3 Source estimation

Source estimation from the cleaned channel-level signals was performed with the FieldTrip toolbox (Oostenveld et al.; 2011) by relying on a source model based on a standard template composed of 15684 uniformly distributed sources in the Montreal Neurological Institute (MNI) space. A non-linear transformation was applied to realign individual EEG sensor positions to the nearest vertex of the scalp mesh (Dykstra et al.; 2012). The geometrical mapping of sources to sensors (namely, the lead-field matrix) was derived by solving the electromagnetic forward problem using a 3-shell boundary element model (BEM) between the vertices of the standard template and the realigned electrodes with the conductivity values of the head tissues set to 0.33 S/m for the skin, 0.0041 S/m for the bone, and 0.33 S/m for the brain. The dimensionality of the obtained lead fields was reduced for each voxel by retaining the source orientation explaining most of the variance. Then, the reduced lead field matrix was used to derive the spatial filter operator by the eLORETA method (Pascual-Marqui et al.; 2011). Finally, the cleaned EEG signals were projected to the source space with the spatial filter matrix, thus obtaining a time-course for each of the 15684 sources.

### 2.4 Connectivity analysis

A seed-based connectivity analysis was performed based on the reconstructed source time-courses. The seed was chosen according to the position of lM1 in the MNI space [–45.9 –9.9 54.6]. The time-courses of the seed and target sources, i.e.; all the other 15683 sources, in the 1-second window prior to the stimulation were band-pass filtered around the individual µ-rhythm frequency, with a bandwidth of 2 Hz, by using a two-pass fourth-order Butterworth filter. The filtered time-courses were padded at both ends by 64 ms and then transformed into their analytic representations by means of the Hilbert transform. Padding was necessary to reduce the edge effects of the filter and the Hilbert transform. Specifically, padding was performed by applying an autoregressive model (Yule-Walker, order 30) in which coefficients were generated from the filtered time-courses. Given the analytic signals of the seed Σ_*S*_(*f, t*) and of each target Σ_*T*_(*f, t*), we extracted the spectral phases *ϕ*_*S*_(*f*) = *arg*{Σ_*S*_(*f*)} and *ϕ*_*T*_(*f*) = *arg*{Σ_*T*_(*f*)} where *arg*{·} represents the argument of a complex-valued number. Finally, the *imaginary part of the phase-locking value* (iPLV) (Palva and Palva, 2012) was estimated as iPLV_*S,T*_(*f*) = | ⟨ ℑ{ exp{*ι* Δ*ϕ*_*S,T*_(*f*)}} ⟩ |, where |·| denotes the absolute value, ℑ{·} is the imaginary part of a complex-valued number, ⟨ · ⟩ indicates expectation value across data epochs and the phase difference Δ*ϕ*_*S,T*_(*f*) was calculated as Δ*ϕ*_*S,T*_(*f*) = *ϕ*_*S*_(*f*) − *ϕ*_*T*_(*f*). We relied on the iPLV metric, because we aimed at characterizing connectivity through phase coupling of neuronal oscillations in line with the CTC hypothesis (Fries, 2015, 2005) with an approach robust to EEG mixing artifacts (Marzetti et al.; 2019).

The procedure described above led to an individual seed-based functional connectivity map at the µ-rhythm frequency. The group averaged functional connectivity map was then computed to identify sources functionally connected to the left M1, in the following termed connectivity Regions of Interest (cROIs), that were considered for subsequent analysis.

Of note, we explicitly decided not to a priori select trials with high signal-to-noise ratio, a procedure employed e.g. in Zrenner et al.; 2018 for phase detection, to avoid potential biases in connectivity analysis or subsequent analyses.

### 2.5 Relation between functional connectivity and Motor Evoked Potential

To assess if the observed functional connectivity is related to the amplitude of the MEP signal at the individual level, for each of the cROIs, we split the trials into two subsets according to the median of the iPLV values: high-connectivity (HC) and low-connectivity (LC) trials. Then, for each cROI and subject, we calculated the MEP change relative to the mean MEP amplitude for the HC and LC trials separately and determined with a paired sample *t*-test whether a difference in these classes of trials exists.

Additionally, we asked whether a set of trials exists for which, taken together, all cROIs exhibit a high or low connectivity to lM1. We termed these subsets as high-connectivity trials for the network (HC_network) and low-connectivity trials for the network (LC_network). The modulation of MEP amplitudes in the HC_network and LC_network trials was assessed, similarly to that between HC and LC trials, by a paired sample *t*-test.

Control analyses were run for the modulation of MEP in different trial subsets defined at subject level according to criteria that do not consider functional connectivity: *i*) splitting the trials into the first and second part of the recording; *ii*) splitting the trials into even and odd. For each of these different trial-splitting approaches, median MEP amplitudes across the first and second subsets were calculated for each subject and a paired sample t-test (two-tails) was run to assess modulations of MEP amplitudes between the two subsets.

### 2.6 Coupling directionality

To assess coupling directionality, we relied on the Multivariate Phase Slope Index (MPSI), known to be more reliable than the corresponding bivariate approaches (Basti et al.; 2018). The calculation of MPSI is based on the estimation of the cross-spectra among time courses of the cROIs and time courses of the voxels surrounding the seed. Specifically, to apply the multivariate directionality metric, we selected all voxels the distance of which is smaller than 4 mm from the seed (subset of dimensionality *n*) and from the cROI centroids. Each multivariate time series was then built as a matrix with the first dimension being *n* and the second being the total data length obtained by concatenating one second prestimulus data for the HC_network and LC_network trials. These time series were used to calculated MPSI over each pair of frequencies in a range spanning 4 Hz centered at the individual µ frequency and with a 1-Hz frequency resolution. To assess the statistical significance of the observed results, we considered a standardized version of MPSI that allows us to interpret the ratio between MPSI and its standard deviation across estimation segments (jackknife approach) as a pseudo-Z score. Finally, a group-level Z-score was obtained by averaging the individual pseudo-Z score values multiplied by the square root of the number of subjects to normalize the variance of the averaged pseudo-Z score distribution. Of note, the coupling directionality measured by MPSI pseudo-Z score cannot be directly interpreted as a measure of the coupling strength, rather it estimates the leader and follower role between a pair of multidimensional signals.

### 2.7 Phase estimation

The phase of µ-rhythm signal at stimulation was estimated by following the approach of (Zrenner et al.; 2018). Specifically, we extracted the signal from the 500 ms preceding the TMS stimulation from the lM1 region in source space, then a forward-backward filter in the individual µ-band (order = 64) was applied and, finally, the filtered signal was trimmed at the beginning and end by 64 ms, to remove the edge effects of the filter. An autoregressive model of order 30 was then used to predict the signal from –64 to +64 ms centered around the TMS stimulation. The phase of the signal was then obtained by applying the Hilbert-transform to the predicted signal and by extracting the phase at time zero. Of note, the phase values and the connectivity values were estimated for all trials that survived artifact rejection. As mentioned in the previous paragraph, no trials were discarded based on EEG power.

### 2.8 Linear regression analysis

To investigate the relation between long-range motor network connectivity, as measured by phase locking (i.e.; phase differences) of µ-rhythm oscillations, and local properties of M1, as measured by the phase of the µ-rhythm oscillation at M1, we tested five different linear models for MEP amplitude prediction at the individual level across all trials.

First, we tested a model in which the connectivity values between lM1 and a single cROI were used as independent variable 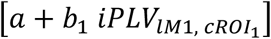; then, we added the connectivity between lM1 and all other (*n* − 1) cROIs to the first model 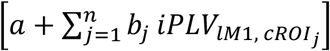. The phase at lM1 was used as the only independent variable in a third model [*a* + *c* cos *φ* + *d* sin *φ*], analogously to Zrenner et al. 2020. Then, all the variables were used in a fourth model including all connectivity and phase as independent variables 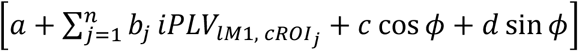. In the above equations *a, b*_1_, *b*_*j*_, *c, d* are the model parameters, *φ* is the phase and 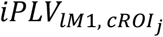 is the connectivity between *ιM*1 and all the cROIs. Finally, a constant model was used as control analysis.

The Akaike Information Criterion (AIC) value (Akaike, 1998) was calculated to compare these different models. Indeed, the AIC-based model selection weights model performance and complexity in a single metric, and the difference between AICs of different models is an indicator of their relative plausibility (Burnham and Anderson, 2004). Specifically, we used AIC to answer the question whether it is worth adding another variable in the model for: connectivity-based model with one cROI versus connectivity-based model with all cROIs; connectivity-based model versus connectivity and phase-based model; phase-based model versus functional connectivity and phase-based model.

## 3. Results

### 3.1 Functional connectivity at the µ-rhythm frequency highlights coupling within the motor network

EEG preprocessing evidenced that, on average, 23% of the channels and 18% of the trials were contaminated by artifacts and were therefore excluded from the following analysis. The average µ-rhythm peak frequency across subjects was 10.5±1.5 Hz. For each subject, a map of iPLV with respect to lM1 was obtained at the individual µ-rhythm peak frequency; the grand average of these individual iPLV maps is shown in Figure 1. The surface-based top view of Figure 1 middle panel shows the location of lM1 (black dot) and the regions functionally connected to it in a color-coded representation in which red indicates high connectivity to lM1. The orthographic views of Figure 1 left and right panels better show the location for the red spots of Figure 1 middle panel and highlight that lM1 is functionally connected to the left Supplementary Motor Area (lSMA, centroid MNI coordinates [–12 –11 74]) and to the right motor cortex (rM1, centroid MNI coordinates [40 –25 52]). While recent EEG reports have detected a correlated pattern of functional connectivity resembling the motor system at broadband (Marino et al.; 2019) and at alpha band (Samogin et al.; 2020) frequencies, current findings indicate the emergence of the motor network at the individual µ-rhythm peak frequency in resting state activity preceding TMS. For following analyses, lSMA and rM1 regions obtained with the described approach were employed as the set of cROIs.

**Figure 1.**
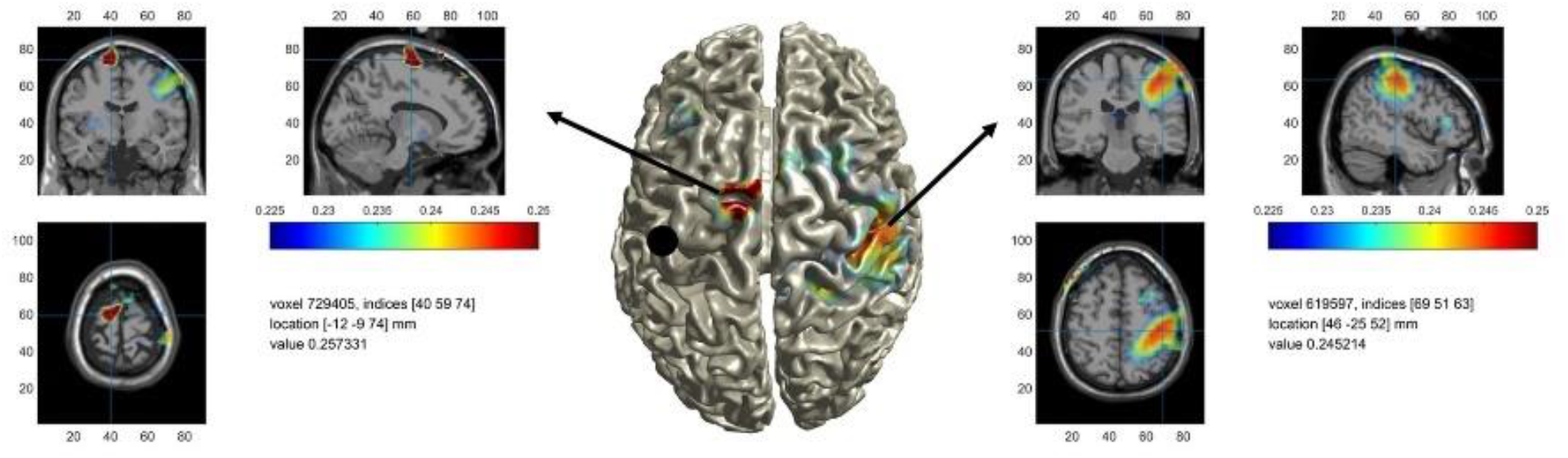
Group averaged functional connectivity (iPLV) to the left primary motor cortex (lM1, black dot in the left hemisphere in the middle panel). The middle panel shows a surface-based projection, and the left and right panels show two different orthographic projections of the same map. All views highlight that the lM1 is functionally connected to the right motor cortex (rM1) and to the left Supplementary Motor Area (lSMA), as defined by the MNI coordinates of their centroids.

### 3.2 MEP amplitude modulates with functional connectivity of the motor network

The modulation of MEP amplitude in the lM1-lSMA HC trials, defined as the percent change of the MEP amplitude with respect to MEP mean value, was calculated for each subject and the average value and its standard error are shown in Figure 2A together with the modulation in lM1-lSMA LC trials. On average, a difference of 21.8 ± 2.5%, (mean ± standard error of the mean) in MEP amplitudes is observed for lM1-lSMA HC trials with respect to lM1-lSMA LC trials (one tail paired-sample *t*-test, *p*=0.03). This result points towards a spontaneous facilitation effect of SMA on M1 at rest, i.e.; a high lM1-lSMA connectivity enhances MEP amplitude, in line with the facilitation obtained by conditioning M1 by stimulating SMA with a cortico-cortical paired associative stimulation protocol (Arai et al.; 2011).

**Figure 2.**
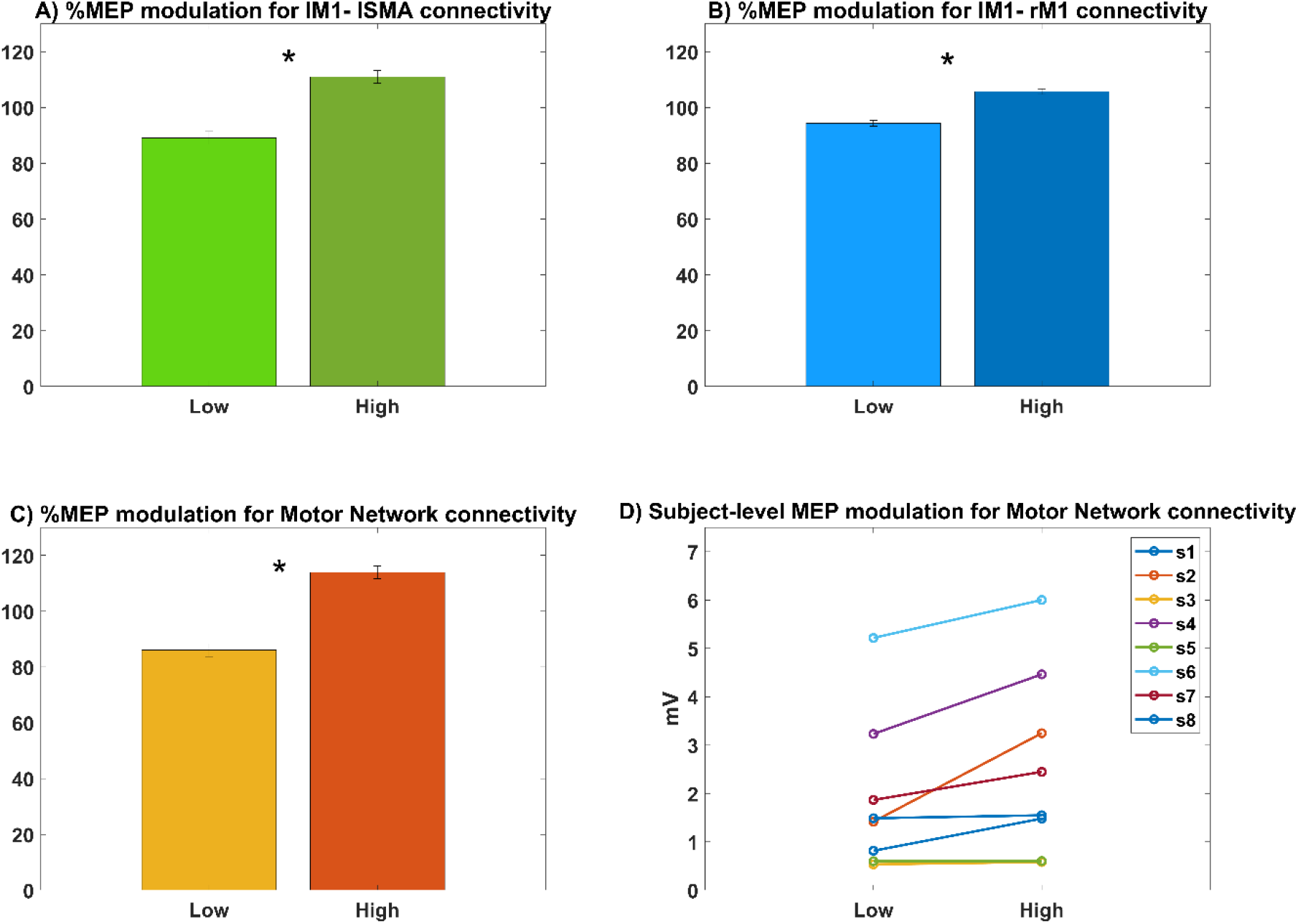
MEP amplitude modulation, percentage values, in trials with high (HC) and low (LC) connectivity, median split based definition. Bars indicate mean MEP amplitude, in the corresponding set of trials, across all subjects and whiskers indicate the standard error of the mean. A) MEP modulation (one-tail paired sample *t*-test, \**p*=0.03) for connectivity between the left motor cortex (lM1) and the left Supplementary Motor Area (lSMA). B) MEP modulation (one-tail paired sample *t*-test, \**p*=0.01) for connectivity between lM1 and the right Motor Cortex (rM1). C) MEP modulation (one-tail paired sample *t*-test, \**p*=0.01) in the subset of trials in which connectivity between lM1 and lSMA and connectivity between lM1 and rM1 are simultaneously low (LC_network) or high (HC_network). D) A positive modulation of connectivity with MEP amplitude (logarithmic value of MEP amplitudes is shown on the y axis) is observed in the majority of the subjects.

Similarly, Figure 2B shows that a modulation of 11.4 ± 1.0% is featured by MEP amplitude in the lM1-rM1 HC trials with respect to lM1-rM1 LC trials (one tail paired-sample t-test, p=0.01). It should be noted that, in general, lM1-lSMA HC trials are a different subset than lM1-rM1 HC trials and similarly for LC trials and that the facilitatory effect of lM1-rM1 connectivity found here is smaller in extent with respect to the lM1-lSMA effect, in line with our observation that the overall functional connectivity between lM1 and rM1 is weaker than that of lM1 with lSMA.

Figure 2C shows the MEP modulation in the subset of trials in which lM1-lSMA and lM1-rM1 functional connectivity trials are both above or both below their corresponding median level. The overlap between HC trials for lM1-lSMA and that for lM1-rM1, across all subjects, is 57% (median value) with an interquartile range of 7%, meaning that the number of trials in which a consistent high or low coupling of the whole network (HC_network or LC_network trials) is observed is above chance level. A significant increase of about 27.9 ± 2.3% for MEP in the HC_network trials is observed with respect to LC_network trials (one tail paired-sample t-test, p=0.01). The network-level MEP modulation is thus increased by about 28% with respect to the largest modulation observed for single node pairs, i.e. lM1-lSMA connectivity.

Finally, Figure 2D shows the modulation of MEP in HC_network and LC_network at the individual level for the eight subjects. While for some of the subjects the positive relation was more evident, other did not show a clear effect. Interestingly, the subjects that show weak or no effect are those which feature low cortico-spinal excitability as indicated by their low MEP amplitude values (subj3, subj5, subj8).

Notably, no significant difference is observable between MEP amplitudes in the first half versus the second half of the trials in the recording (paired-sample *t*-test, *p*=0.20). This analysis allowed us to rule out possible habituation or potentiation effects induced by the high number of stimuli delivered in each experimental session. Similarly, MEP amplitudes in even versus odd trials were not significantly different (paired-sample *t*-test, *p*=0.88), thus allowing to exclude chance effects induced by the trial splitting procedure. Finally, we calculated functional connectivities between lM1 and lSMA and between lM1 and rM1 and their relation to MEP amplitudes for the theta (average value across subjects: 5.0 ± 0.7 Hz) and beta (average value across subjects: 21 ± 3 Hz) frequencies. These analyses revealed no significant effect of functional connectivity between lM1 and lSMA or between lM1 and rM1 on MEP amplitude modulation for theta (lM1-lSMA: one tail paired-sample *t*-test, *p*=0.10; lM1-rM1: one tail paired-sample *t*-test, *p*=0.23) and beta (lM1-lSMA: one tail paired-sample *t*-test, *p*=0.47; lM1-rM1: one tail paired-sample *t*-test, *p*=0.09).

### 3.3 Coupling directionality reveals a top-down control of SMA on bilateral M1

The MPSI analysis revealed that in the high functional connectivity trials, coupling directionality, averaged across subjects, indicates a connectivity from lSMA to lM1 (pseudo-Z = –3.61, *p*=3*10^-4^) and to rM1 (pseudo-Z = –2.53, *p*=0.01). Conversely, no significant directionality could be assessed for the connectivity between lM1 and rM1 (pseudo-Z = –1.88, *p*=0.06). Similar results were obtained for low connectivity trials (lSMA-lM1 pseudo-Z = –4.78 *p*=2*10^-6^, lSMA-rM1 pseudo-Z = –2.48 *p*=0.01, lM1-rM1 pseudo-Z = –1.21 *p*=0.23).

### 3.4 A linear regression model that relies on network connectivity and lM1 phase best predicts MEP

The results for the comparison between a linear regression model, at single subject level, in which the MEP amplitude is predicted only by the functional connectivity between lM1 and lSMA and a model in which functional connectivity of lM1-rM1 is added as second independent variable (i.e.; Motor Network model) are reported in Supplementary Material Table SM1 and Table SM2. These data indicate that the Motor Network model performs overall better than the lM1-lSMA model. In the following, we will thus further compare, in terms of their respective AIC values, the Motor Network model with different linear regression models in which MEP amplitude is predicted only by the phase at lM1 (Table 1, column 3), a model in which MEP is predicted by network-level functional connectivity (Table 1, column 4), and a model in which both are used as independent variables to predict MEP (Table 1, column 5). Additionally, AIC of MEP prediction by a constant model is reported (Table 1, column 2). Column 6 of Table 1 indicates the model to be preferred according to the criteria defined in (Burnham and Anderson, 2004).

**Table 1.**
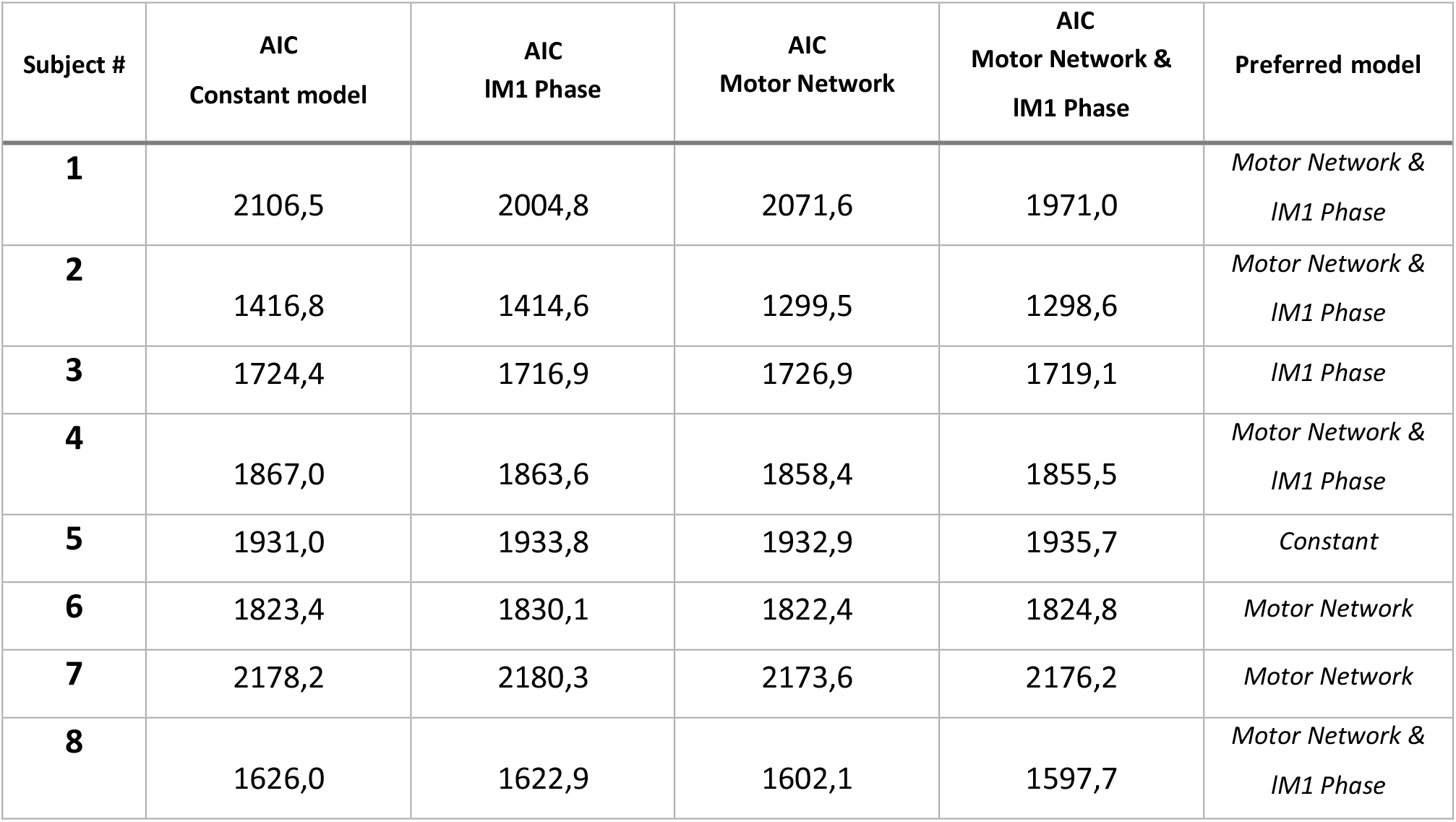

Overall, the model with Motor Network and lM1 Phase as independent variables was preferred in 4 out of 8 subjects, while a model with Motor Network only as independent variable is preferred in 2 out of 8 subjects. In the remaining 2 subjects, either the Phase only model or the constant model are preferred.

Table 2 shows the plausibility of all the tested models.

**Table 2.**

| Subject # | Constant model | IM1 Phase | Motor Network | Motor Network & IM1 Phase |
| --- | --- | --- | --- | --- |
| **1** | Not plausible | Not plausible | Not plausible | Preferred |
| **2** | Not Plausible | Not Plausible | Plausible | Preferred |
| **3** | Not plausible | (Mildly)<br>Preferred | Not plausible | Plausible |
| **4** | Not Plausible | Not Plausible | Plausible | Preferred |
| **5** | Preferred | Plausible | Plausible | Plausible |
| **6** | Plausible | Not Plausible | Preferred | Plausible |
| **7** | Not Plausible | Not Plausible | Preferred | Plausible |
| **8** | Not plausible | Not plausible | Not plausible | Preferred |

Overall, tables 1 and 2 indicate that including local (lM1 phase) and long-range (Motor Network) characteristics of the source-space EEG signal results in the best or in a plausible predictive model for single trial MEP amplitudes.

## 4. Discussion

In the present proof-of-concept study, we show that EEG-derived long-range connectivity of the primary motor cortex (M1) in the pre-stimulation period at individual µ-rhythm peak frequency is largely congruent with the motor network and that the connectivity state of this network modulates the motor responses evoked by transcranial magnetic stimulation of the primary motor cortex. Specifically, stronger coupling of left M1 with left supplementary motor area (SMA) and right M1, as measured by phase locking of µ-rhythm oscillations, was related to larger motor evoked potential (MEP) amplitudes and vice versa. These findings indicate that the corticospinal excitability is associated with the coordinated interaction among key areas of the motor network rather than only with the local activity of M1. Importantly, the observed positive relation between motor network connectivity and MEP amplitudes holds at the individual level, even if not for all subjects. Subjects that do not show the effect feature an overall low cortico-spinal excitability, as indexed by low MEP values, across the whole recording. Thus, we speculate that lack of modulation is due to a generally low responsiveness and not to the relation to motor network connectivity.

Previous work investigating an association between MEP amplitude and pre-stimulus EEG phase-locking, as measured by coherence magnitude, has observed a coupling between the stimulated primary motor cortex and a large swath of centro-parietal cortex in the delta band and of frontal cortex in the high beta band (Ferreri et al.; 2014). Yet, the present findings indicate that brain connectivity states affect the corticospinal excitability in a topographically selective (i.e.; motor network) fashion. Our result was likely obtained by using a source level functional connectivity approach based on a metric robust to field spread and volume conduction effects (Marzetti et al.; 2019), as compared to the approach used in (Ferreri et al.; 2014). The more recent study from Vetter et al. (2021) investigates, in a real time EEG-TMS experiment, the association between MEP amplitude and pre-stimulus EEG phase-locking between two specific EEG channels (after application of a Hjorth montage) located approximately above the motor cortices. Overall, the study concludes that functional connectivity was predictive of cortico-spinal excitability together with power and phase. Nevertheless, this study employs seed-based sensor-level connectivity analysis which makes it impossible to assess whether the considered signals actually come from motor areas. Similarly, the real-time setting available for the experiment did not allow to consider more than two channels and thus to investigate the potential augmentation of the observed effect when more than just two regions in a network are connected.

The positive relationship between MEP amplitude and pre-stimulus motor network connectivity is consistent with neurophysiological and neuroimaging lines of evidence from functional Magnetic Resonance Imaging indicating that such functional interactions are relevant for MEP (Cárdenas-Morales et al.; 2014) and for hand function in healthy individuals (Pool et al.; 2013). Clinically, it has been observed that intra-hemispheric M1-SMA (Grefkes et al.; 2010; Liu et al.; 2021) and inter-hemispheric M1-M1 (Carter et al.; 2010; Grefkes et al.; 2010) functional connections are behaviorally relevant for recovery after motor stroke as well as predictive of functional improvement induced by theta-burst TMS stimulation (Diekhoff-Krebs et al.; 2017). Moreover, our results for coupling directionality within the motor network indicate an overall stable intrinsic coupling directionality from lSMA to lM1 and rM1, in line with previous evidence of SMA conditioning effect on M1 excitability (Arai et al.; 2011) as well as with dynamic causal modelling-based SMA bilateral control over primary motor cortices (Pool et al.; 2013).

In addition, the lack of a dominant directionality between the two motor cortices in the resting brain is in line with Grefkes et al. (2008).

Furthermore, our individual analysis results show that, for the majority of the subjects, a model including both long-range phase locking within the motor network and the phase of M1 oscillation results in a better prediction of MEP amplitudes than either one of these factors alone. A model based on motor network connectivity alone is, in general, more plausible than a model based on M1 phase alone. Of note, in contrast to previous works investigating the role of phase in predicting MEP amplitudes, e.g. (Zrenner et al.; 2018), we did not select trials on a power-based criterion (Zrenner et al.; 2020); this might justify the difference between our results and previous ones for phase-based prediction.

Finally, our findings pair to the work by Stefanou et al. (Stefanou et al.; 2018) in supporting the idea that functional connectivity can be directly exploited to design paired- or multi-coil stimulation protocols in which the stimulation is delivered when nodes in the network are phase-coupled. Indeed, the functional connectivity approach used in our paper can be extended to real-time estimation (Sommariva et al.; 2019; Basti et al.; 2022) and, thus, translated into protocols for state-dependent connectivity-based stimulation.

Although we acknowledge that a limitation of this study is in the limited size of our cohort, it must be noted that our data rely on a high number of trials for each subject (overall number of trails about 8000). The final aim of our study is to provide a proof-of-concept of the possibility to extract an EEG-based motor network from EEG-TMS data, as well as to assess the relation between motor network connectivity and cortico-spinal excitability at single subject level and to possibly take advantage of our results for an individualized connectivity targeted stimulation. For this reason, relying on many trials per subjects allows us to perform such an investigation in a robust manner in this study in which a subset of trials with high or low functional connectivity must be considered to these purposes.

## 5. Conclusions

To the best of our knowledge, this is the first study that investigates to what extent the connectivity state of a brain network in source space prior to transcranial magnetic stimulation influences its outcome. Here, we specifically addressed this question for the network linked to the left primary motor cortex at the individual peak frequency of the sensorimotor µ-rhythm. We demonstrate that a high-connectivity state within this network, which largely overlaps with the motor network topography, features a facilitation effect on the amplitude of the motor evoked potential induced by left primary motor cortex stimulation. Notably, the increase of MEP amplitude with enhanced motor network connectivity supports the idea that connectivity-informed real-time state-dependent stimulation may have a high potential including a therapeutic efficacy.

## Conflicts of interest

The authors declare the following financial interests/personal relationships which may be considered as potential competing interests: R.J.I. has received consulting fees from Nexstim plc and has patents and patent applications on TMS technology. C.Z. reports an interest in a start-up company, sync2brain GmbH, which commercializes the real-time EEG analysis technology. U.Z. received grants from the German Ministry of Education and Research (BMBF), German Research Foundation (DFG), Takeda Pharmaceutical Company Ltd.; and consulting fees from CorTec GmbH, all not related to this work. The other authors declare that they have no known competing financial interests or personal relationships that could have appeared to influence the work reported in this paper.

## Credit authorship contribution statement

**Laura Marzetti**: Conceptualization, Methodology, Software, Writing - Original Draft, Writing - Review & Editing; **Alessio Basti**: Software, Writing - Review & Editing; **Antonello Baldassarre**: Writing - Original Draft, Writing - Review & Editing; **Roberto Guidotti**: Software, Writing - Review & Editing; **Johanna Metsomaa**: Data Curation, Writing - Review & Editing; **Christoph Zrenner**: Investigation, Writing - Review & Editing; **Antea D’Andrea:** Writing - Review & Editing; **Saeed Makkinayeri:** Writing - Review & Editing; **Giulia Pieramico:** Writing - Review & Editing; **Risto Ilmoniemi**: Writing - Review & Editing, Funding acquisition; **Ulf Ziemann**: Writing - Review & Editing, Funding acquisition; **Gian Luca Romani**: Writing - Review & Editing, Funding acquisition; **Vittorio Pizzella**: Conceptualization, Methodology, Writing - Original Draft, Writing - Review & Editing.

## Acknowledgements

The authors wish to thank Filippo Zappasodi and Timo Roine for useful discussions on the work presented in this manuscript, as well as Maria Ermolova for data sharing.

This study was funded by the European Research Council (ERC Synergy) under the European Union’s Horizon 2020 research and innovation programme (ConnectToBrain; grant agreement No. 810377). The content of this article reflects only the author’s view and the ERC Executive Agency is not responsible for the content.

V.P. acknowledges financial support under the National Recovery and Resilience Plan (NRRP), Mission 4, Component 2, Investment 1.1, Call for tender No. 104 published on 2.2.2022 by the Italian Ministry of University and Research (MUR), funded by the European Union – NextGenerationEU– Project Title “Adaptive Brain Connectivity and Cognition ABC&C” – Grant Assignment Decree No. 1349 adopted on 25.08.2023 by the Italian Ministry of Ministry of University and Research (MUR). L.M. acknowledges financial support under the National Recovery and Resilience Plan (NRRP), Mission 4, Component 2, Investment 1.1, Call for tender No. 1409 published on 14.9.2022 by the Italian Ministry of University and Research (MUR), funded by the European Union – NextGenerationEU– Project Title “NEUROSTAR BTP – NEUROplasticity STimulation to Ameliorate Results of Surgery in Brain Tumor Patients”-Grant Assignment Decree No. 1367 adopted on 01.09.2023 by the Italian Ministry of Ministry of University and Research (MUR) and from the Italian Ministry of Ministry of University and Research (MUR) National Operational Programme Research and Innovation 2014-2020 additional funds ESF-REACT-EU (Decree No. 1062 adopted on 08.10.2021 by MUR).

## Notes

### Summary of Updates

Double-checked statistical results, modified figure 2, improved text, modified title

