## Supplementary Material for "Exploring Motor Network Connectivity in State-Dependent Transcranial Magnetic Stimulation: A Proof-of-Concept Study"

Table SM1 shows the results, at single subject level, for the comparison between a linear regression model in which the MEP amplitude is predicted only by the functional connectivity between lM1 and lSMA and a model in which functional connectivity of lM1-rM1 is added as second independent variable. Columns 2 and 3 report the AIC values for the two models, while column 4 indicates absolute value of the AIC difference between the model with the largest AIC and the model with the smallest AIC for each subject. As a rule of thumb, the larger the AIC difference between the two models the more the evidence against the model with the larger AIC. The corresponding model plausibility, according to the criteria defined in Burnham and Anderson (2002), is shown in column 5.

**Table SM1**

| Subject # | AIC<br>lM1-lSMA model | AIC<br>Motor Network model | ∣AIC difference∣ | Preferred model |
|---|---|---|---|---|
| **1** | 2069.5 | 2071.6 | 2.1 | lM1-lSMA model |
| **2** | 1315.0 | 1299.5 | 15.5 | Motor network model |
| **3** | 1726.5 | 1726.9 | 0.4 | No preference |
| **4** | 1858.7 | 1858.4 | 0.3 | No preference |
| **5** | 1933.2 | 1932.9 | 0.3 | No preference |
| **6** | 1825.8 | 1822.5 | 3.3 | No preference |
| **7** | 2171.7 | 2173.7 | 2.0 | Motor network model |
| **8** | 1613.3 | 1602.1 | 11.2 | No preference |

Overall, in the 25% of the subjects, the motor network model is preferred over the model that considers lM1-lSMA connectivity values only, consistently with the network-level facilitatory effect described above, while the latter is preferred with respect to the former only in the 12.5% of the subjects. In the remaining 62.5% of the subjects, no preference between the two models was observed.

Table SM2 shows the plausibility of the two models and indicates that the Motor Network model performs overall better than the lM1-lSMA model.

**Table SM2**

| Subject # | lM1-lSMA model | Motor Network model |
|---|---|---|
| **1** | Preferred | Plausible |
| **2** | Not Plausible | Preferred |
| **3** | Preferred | Plausible |
| **4** | Plausible | Preferred |
| **5** | Plausible | Preferred |
| **6** | Plausible | Preferred |
| **7** | Preferred | Plausible |
| **8** | Not Plausible | Preferred |
